# Learning interpretable cellular responses to complex perturbations in high-throughput screens

**DOI:** 10.1101/2021.04.14.439903

**Authors:** Mohammad Lotfollahi, Anna Klimovskaia Susmelj, Carlo De Donno, Yuge Ji, Ignacio L. Ibarra, F. Alexander Wolf, Nafissa Yakubova, Fabian J. Theis, David Lopez-Paz

**Author notes:** These authors contributed equally to the work (Mohammad Lotfollahi, Anna Klimovskaia Susmelj). These authors contributed equally to the work (Carlo De Donno, Yuge Ji). Present address: Cellarity, Inc., Cambridge, MA.

## Abstract

Recent advances in multiplexed single-cell transcriptomics experiments are facilitating the high-throughput study of drug and genetic perturbations. However, an exhaustive exploration of the combinatorial perturbation space is experimentally unfeasible, so computational methods are needed to predict, interpret, and prioritize perturbations. Here, we present the compositional perturbation autoencoder (CPA), which combines the interpretability of linear models with the flexibility of deep-learning approaches for single-cell response modeling. CPA encodes and learns transcriptional drug responses across different cell type, dose, and drug combinations. The model produces easy-to-interpret embeddings for drugs and cell types, which enables drug similarity analysis and predictions for unseen dosage and drug combinations. We show that CPA accurately models single-cell perturbations across compounds, doses, species, and time. We further demonstrate that CPA predicts combinatorial genetic interactions of several types, implying that it captures features that distinguish different interaction programs. Finally, we demonstrate that CPA can generate *in-silico* 5,329 missing genetic combination perturbations (97.6% of all possibilities) with diverse genetic interactions. We envision our model will facilitate efficient experimental design and hypothesis generation by enabling *in-silico* response prediction at the single-cell level, and thus accelerate therapeutic applications using single-cell technologies.

## Introduction

Single-cell RNA-sequencing (scRNA-seq) profiles gene expression in millions of cells across tissues[1, 2] and species[3]. Recently, novel technologies have been developed that extend these measurements to high-throughput screens (HTSs), which measure response to thousands of independent perturbations[4, 5]. These advances show promise for facilitating and thus accelerating drug development[6]. HTSs applied at the single-cell level provide both comprehensive molecular phenotyping and capture heterogeneous responses, which otherwise could not be identified using traditional HTSs[4].

While the development of high-throughput approaches such as “cellular hashing” [4, 7, 8] facilitates scRNA-seq in multi-sample experiments at low cost, these strategies require expensive library preparation[4], and do not easily scale to large numbers of perturbations. These shortcomings become more apparent when exploring the effects of combination therapies[9–11] or genetic perturbations[5, 12, 13], where experimental screening of all possible combinations becomes infeasible. While projects such as the Human Cell Atlas[14] aim to comprehensively map cellular states across tissues in a reproducible fashion, the construction of a similar atlas for the effects of perturbations on gene expression is impossible, due to the vast number of possibilities. Since brute-force exploration of the combinatorial search space is infeasible, it is necessary to develop computational tools to guide the exploration of the combinatorial perturbation space to nominate promising candidate combination therapies in HTSs. A successful computational method for the navigation of the combinatorial space must be able to predict the behaviour of cells when subject to novel combinations of perturbations only measured separately in the original experiment. These data are referred to as Out-Of-Distribution (OOD) data. OOD prediction would enable the study of perturbations in the presence of different treatment doses [4, 15], combination therapies[8], multiple genetic knockouts[5], and changes across time[15].

Recently, several computational approaches have been developed for predicting cellular responses to perturbations[16–20]. The first approach leverages mechanistic modeling [18, 19] to predict cell viability[19] or the abundance of a few selected proteins[18]. Although they are powerful at interpreting interactions, mechanistic models usually require longitudinal data (which is often unavailable in practice) and most do not scale to genome wide measurements to predict high-dimensional scRNA-seq data. Linear models[12, 21] do not suffer from these scalability issues, but have limited predictive power and are unable to capture nonlinear cell-type specific responses. In contrast, deep learning (DL) models do not face these limitations. Recently, DL methods have been used to model gene expression latent spaces from scRNA-seq data [22–25], and describe and predict single-cell responses [16, 17, 20, 26]. However, current DL-based approaches also have limitations: they model only a handful of perturbations; can be difficult to interpret; cannot handle combinatorial treatments; and cannot incorporate continuous covariates such as dose and time, or discrete covariates such as cell types, species, and patients while preserving interpretability. Therefore, while current DL methods have modeled individual perturbations, none have been proposed for HTS.

Here, we propose the *compositional perturbation autoencoder (CPA)*, a novel, interpretable method to analyze and predict scRNA-seq perturbation responses across combinations of conditions such as dosage, time, drug, and genetic knock-out. The CPA borrows ideas from interpretable linear models, and applies them in a flexible DL model to learn factorized latent representations of both perturbations and covariates. Given a scRNA-seq dataset, the perturbations applied, and covariates describing the experimental setting, CPA decomposes the data into a collection of embeddings (representations) associated with the cell type, perturbation, and other external covariates. Since these embeddings encode the transcriptomic effect of a drug or genetic perturbation, they can be used by CPA users to study drug effects and similarities useful for drug repurposing applications. By virtue of an adversarial loss, these embeddings are independent from each other, so they can be recombined at prediction time to predict the effect of novel perturbation and covariate combinations. Therefore, by exploring novel combinations, CPA can guide experimental design by directing hypotheses towards expression patterns of interest to experimentalists. We demonstrate the usefulness of CPA on five public datasets and multiple tasks, including the prediction and analysis of responses to compounds, doses, time-series information, and genetic perturbations.

## Results

### Multiple perturbations as compositional processes in gene expression latent space

Prior work has modeled the effects of perturbations on gene expression as separate processes. While differential expression compares each condition separately with a control, modeling a joint latent space with a conditional variational autoencoder[17, 26, 27] is highly uninterpretable and not amenable to the prediction of the effects of combinations of conditions. Our goal here is to factorize the latent space of neural networks to turn them into interpretable, compositional models. If the latent space were linear, we could describe the observed gene expression as a factor model where each component is a single perturbation.

However, gene expression latent spaces, particularly in complex tissues, are nonlinear and best described by a graph or nonlinear embedding approximations[28, 29]. In scRNA-seq datasets, gene expression profiles of cell populations are often observed under multiple perturbations such as drugs, genetic knockouts, or disease states, in labeled covariates such as cell line, patient, or species. Each cell is labeled with its experimental condition and perturbation, where experimental covariates are captured in categorical labels and perturbations are captured using a continuous value (e.g. a drug applied with different doses). This assumes a sufficient number of cells per condition to permit the estimation of the latent space in control and perturbation states using a large neural network.

Instead of assuming a factor model in gene expression space, we instead model the nonlinear superposition of perturbation effects in the nonlinear latent space, in which we constrain the superposition to be additive (see **Methods**). We decouple the effects of perturbations and covariates, and allow for continuous effects such as drug dose by encoding this information in a nonlinearly transformed scalar weight: a learned drug-response curve. The linear latent space factor model enables interpretation of this space by disentangling latent space variance driven by covariates from those stemming from each perturbation. At evaluation time, we are able to not only interpolate and interpret the observed perturbation combinations, but also to predict other combinations, potentially in different covariate settings.

### Compositional perturbation autoencoder (CPA)

We introduce the CPA (see **Methods**), a method combining ideas from natural language processing [30] and computer vision [31, 32] to predict the effects of combinations of perturbations on single-cell gene expression. Given a single-cell dataset of multiple perturbations and covariates, the CPA first uses an encoder neural network to decompose the cells’ gene expression into three learnable, additive embeddings, which correspond to its basal state, the observed perturbation, and the observed covariates. Crucially, the embedding that the CPA encoder learns about a cell’s basal state is disentangled from (does not contain information about) the embeddings corresponding to the perturbation and the covariates. This disentangling is achieved by training a discriminator classifier in a competition against the encoder network of the CPA. The goal of the encoder network in the CPA is to learn an embedding representing a cell’s basal state, from which the discriminator network cannot predict the perturbation or covariate values. To perform well, the embedding of the cell’s basal state should contain all of the information about the cell’s specifics. To account for continuous time or dose effects, the learned embeddings about each perturbation are scaled nonlinearly via a neural network which receives the continuous covariate values for each cell, such as the time or the dose. After integration of the learned embeddings about the cell’s basal state, perturbations, and covariate values into an unified embedding, the CPA uses a neural network decoder to recover the cell’s gene expression vector (**Figure 1**). Similar to many neural network models, the CPA is trained using backpropagation [33] on the reconstruction and discriminator errors (see **Methods**), to tune the parameters of the encoder network, the decoder network, the embeddings corresponding to each perturbation and covariate value, and the dose/time nonlinear scalers. The learned embeddings allow the measurement of similarities between different perturbations and covariates, in terms of their effects on gene expression. The main feature of the CPA is its flexibility of use at evaluation time. After obtaining the disentangled embeddings corresponding to some observed gene expression, perturbation, and covariate values, we can intervene and swap the perturbation embedding with any other perturbation embedding of our choice. This manipulation is effectively a way of estimating the answer to the counterfactual question: what would the gene expression of this cell have looked like, had it been treated differently? This approach is of particular interest in the prediction of unseen perturbation combinations and their effects on gene expression. The CPA can also visualize the transcriptional similarity and uncertainty associated with perturbations and covariates, as later demonstrated.

**Figure 1:**
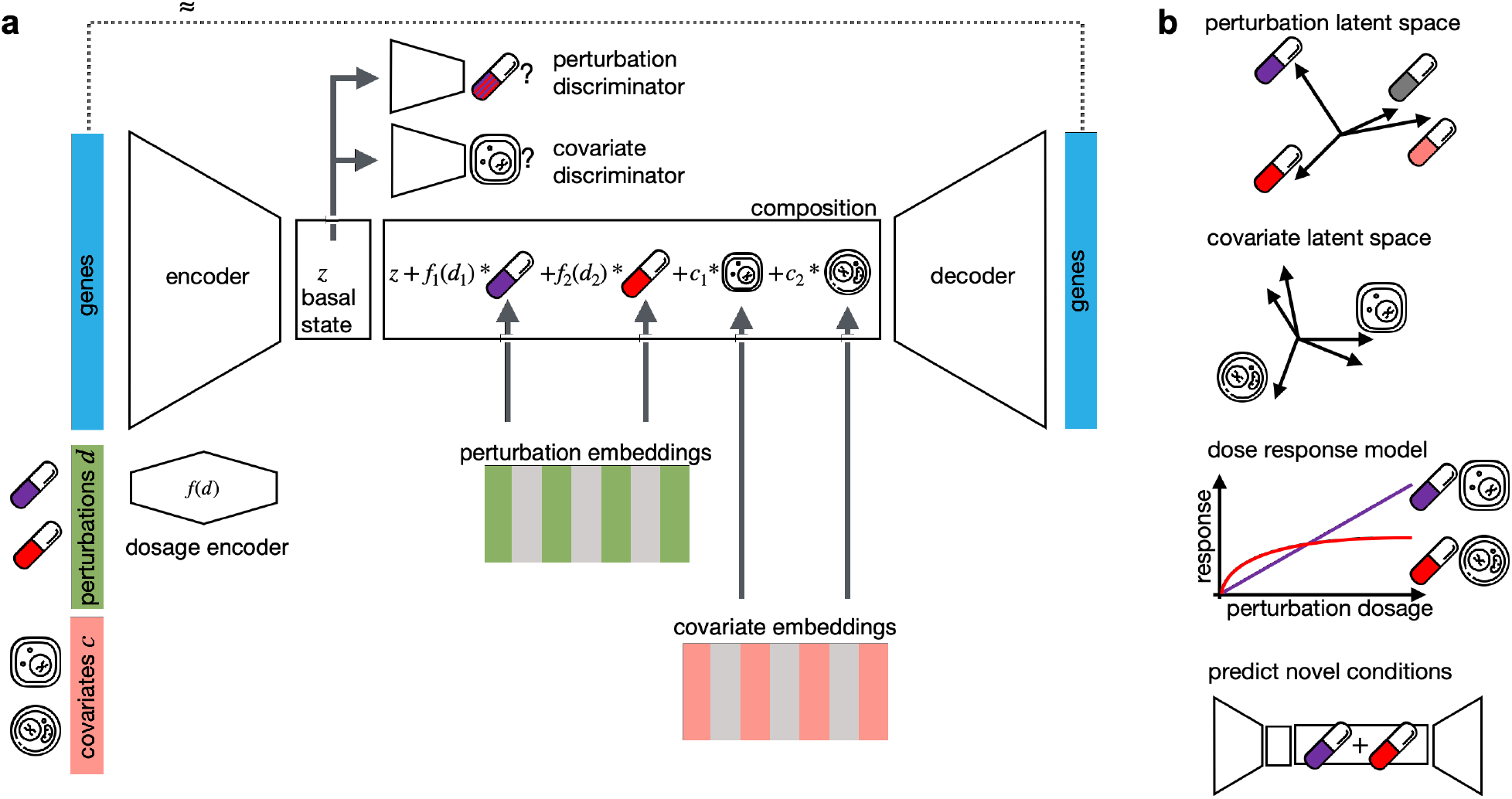
Interpretable single-cell perturbation modeling using a compositional perturbation autoencoder (CPA). **(a)** Given a matrix of gene expression per cell together with annotated potentially quantitative perturbations *d* and other covariates such as cell line, patient or species, CPA learns the combined perturbation response for a single-cell. It encodes gene expression using a neural network into a lower dimensional latent space that is eventually decoded back to an approximate gene expression matrix, as close as possible to the original one. To make the latent space interpretable in terms of perturbation and covariates, the encoded gene expression vector is first mapped to a “basal state” by feeding the signal to discriminators to remove any signal from perturbations and covariates. The basal state is then composed with perturbations and covariates, with potentially reweighted dosages, to reconstruct the gene expression. All encoder, decoder and discriminator weights as well as the perturbation and covariate dictionaries are learned during training. **(b)** Features of CPA are interpreted via plotting of the two learned dictionaries, interpolating covariate-specific dose response curves and predicting novel unseen drug combinations.

### CPA allows predictive and exploratory analyses of single-cell perturbation experiments

We first demonstrated the performance and functionality of the CPA on three small single-cell datasets (**Figure 2**): a Sci-Plex2 dataset of human lung cancer cells perturbed by four drugs [35], a 96-plex-scRNA-seq experiment of HEK293T under different drug combinations [8], and a longitoudnal cross-species dataset of lipopolysaccharide (LPS) treated phagocytes [15] (see **Methods**).

**Figure 2:**
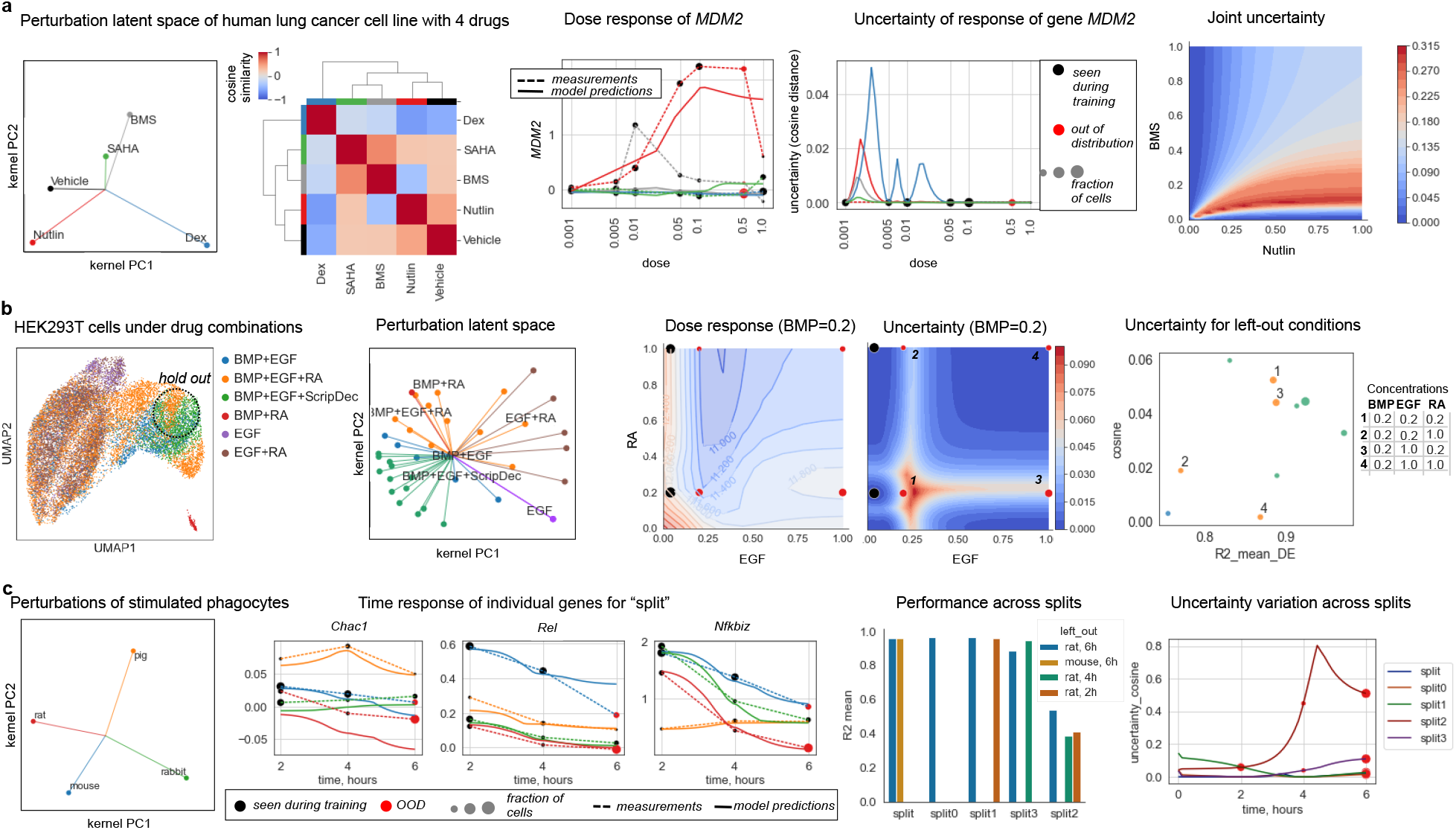
The CPA learns an interpretable latent space learning across drug dosages, drug combinations and experimental systems. **(a)** The sci-Plex 2 dataset from Srivatsan et al [34]. Dose-response curves were generated using the CPA as a transfer from Vehicle cells to a given drug-dose combination. The *MDM2* gene, the top gene differentially expressed after treatment with Nutlin, was selected as an example. Black dots on the dose-response curve denote points seen at training time, red dots denote examples held out for OOD predictions. The sizes of the dots are proportional to the number of cells observed in the experiment. Solid lines correspond to the model predictions, dashed lines correspond to the linear interpolation between measured points. Nutlin and BMS are selected as examples of uncertainty in predictions for drug combinations. **(b)** 96-plex-scRNA-seq experiment from Gehring et al [8], with UMAP, showing variation of responses in gene expression space. The dashed circle on the UMAP represents the area on the UMAP where the majority of the cells from the left-out (OOD) condition lie. The experiment did not contain samples of individual drugs; therefore we represented the latent space of the drug combinations measured in the experiment. The dose-response surface was obtained via model predictions for a triplet of drugs: BMS at a fixed dose of 0.2, and EGF and RA changing on a grid. **(c)** Cross-species dataset from Hagai et al [15], with samples of rat and mouse at time point 6 held out from training, and used as OOD. The latent space representation of individual species, and the individual average response of a species across time, demonstrates that the species are fairly different, with a small similarity between rat and mouse. The time response curves of individual genes demonstrate that the model is able to capture nonlinear behavior. The OOD splits benchmark demonstrates the way in which model performance on the distribution case changes when the model is trained on different subsets of the data. Split2 corresponds to the most difficult case, where all three time points for rat were held out from training. Red dots denote examples held out for OOD predictions; the size is of the dots is proportional to the number of cells observed in the experiment.

All three datasets represent different scenarios of the model application: (i) diverse doses; (ii) drug combinations; and (iii) several Species and variation with respect to time instead of dose. We split each dataset into three groups: train (used for model training), test (used for tuning of the model parameters), and OOD (never seen during training or parameter setting, and intended to measure the generalization properties of the model). **Supplementary Tables** 1–5 shows the *R*^2^ metrics (see Methods) for the performance of the CPA on these datasets and various splits.

Sci-plex from Srivatsan et al. [35] contains measurements of a human lung adenocarcinoma cell line treated with four drug perturbations at increasing doses. In this scenario, the model learns to generalize to the unseen dosages of the drugs. To demonstrate the OOD properties, we withheld cells exposed to the second to largest dose among all drugs. This choice was made because the vast majority of cells are dead for most of the drugs at the highest dosage, and we would not have enough cells to statistically test the generalizability of the CPA model. Since the latent space representation learned by the CPA is still high-dimensional, we can use various dimensionality reduction methods to visualize it, or simply depict it as a similarity matrix (**Figure 2a**). In **Supplementary Table 2** we compare the performance of the CPA on the OOD example on two simple baselines: taking the maximum dose as a proxy to the previous dose, and a linear interpolation between two measured doses. These results demonstrate that the model consistently achieves high scores (a maximum score of 1 yields perfect reconstruction) on all of the OOD cases, and on two of them significantly outperform the baselines for Nutlin (0.92 vs 0.85) and BMS (0.94 vs 0.89). To demonstrate how well the CPA captured the dose-response dynamics of individual genes, we looked at the top differentially expressed genes upon Nutlin perturbation (**Figure 2a**). The dose-response curve agrees well with the observed data. We additionally propose a simple heuristic to measure the model’s uncertainty (see **Methods**) with respect to unseen perturbation conditions. The model shows very low uncertainty on the OOD split. This observation agrees well with the CPA’s high *R*^2^ scores on the OOD example. However, when we tested the uncertainty of the model on a combination of two drugs (**Figure 2a**), we saw that it produces much higher uncertainty compared to single drugs. This finding agrees with the fact that the model never saw some drug combinations during training, and that such predictions are more unreliable.

As a the second working example, we took the 96-plex-scRNAse dataset from Gehring et al. [8]. This dataset contains 96 unique growth conditions using combinations of various doses of four drugs applied to HEK293T cells. We hold out several combinations of these conditions as OOD cases, as detailed in (**Supplementary Table 3**). We show that the CPA is able to reliably predict expression patterns of unseen drug combinations (**Supplementary Table 3**) and produce a meaningful latent perturbation latent space (**Figure 2b**). For this dataset, even simple baselines are not applicable anymore, since the expression of cells exposed to the individual drugs were not measured. We also confirmed that our heuristic for the measurement of uncertainty agreed with the model’s performance on OOD examples.

As our third example we studied the cross-species dataset from Hagai *et al.*[15]. Here we show that the CPA can also be applied in the setting of multiple covariates, such as different species or cell types, and the dynamics of the covariate can be a non-monotonic function, such as time instead of the dose-response. In this example, bone marrow-derived mononuclear phagocytes from mouse, rat, rabbit, and pig were challenged with LPS (**Figure 2c**). The learned CPA latent space agreed with expected species similarities, with a relatively higher value found between rat and mouse. We compared the generalization abilities of the model by withholding different parts of the data for OOD cases: “splitO” (rat at six hours), “split1” (rat at two and six hours), “split2” (rat at two, four, and six hours), “split3” (rat at four and six hours), and “split” (rat and mouse at six hours. This last split was used for the main analysis) (**Supplementary Table 4**). The model produced high performance values compared to the performance on the test split (see **Supplementary Table 5**) on the majority of the OOD splits, and showed a comparatively lower performance when the model was not exposed to any LPS and rat examples with the exception of control cells. On this dataset, we observed that the model with the lowest performance was the one with the highest number of held-out examples, yet the model uncertainty also spiked for these OOD cases, suggesting that they might be not reliable (**Figure 2c**). In contrast, for cases with high *R*^2^ scores, models were more certain about these predictions (**Supplementary Table 4**).

### CPA finds interpretable latent spaces in large-scale single-cell high-throughput screens

The recently proposed sci-Plex assay [35] profiles thousands of independent perturbations in a single experiment via nuclear hashing. With this high-throughput screen, 188 compounds were tested in 3 cancer cell lines. The panel was chosen to target a diverse range of targets and molecular pathways, covering transcriptional and epigenetic regulators and diverse mechanisms of action. The screened cell lines A549 (lung adenocarcinoma), K562 (chronic myelogenous leukemia), and MCF7 (mammary adenocarcinoma) were exposed to each of these 188 compounds at four doses (10 nM, 100 nM, 1 *μ*M, 10 *μ*M), and scRNA-seq profiles were generated for altogether 290 thousand cells (**Figure 3a**). As above, we split the dataset into 3 subsets: train, test, and OOD. For the OOD case, we held out the highest dose (10 *μ*M) of the 36 drugs with the strongest effect in all three cell lines. Drug, dose, and cell line combinations present in the OOD cases were removed from the train and test sets.

**Figure 3:**
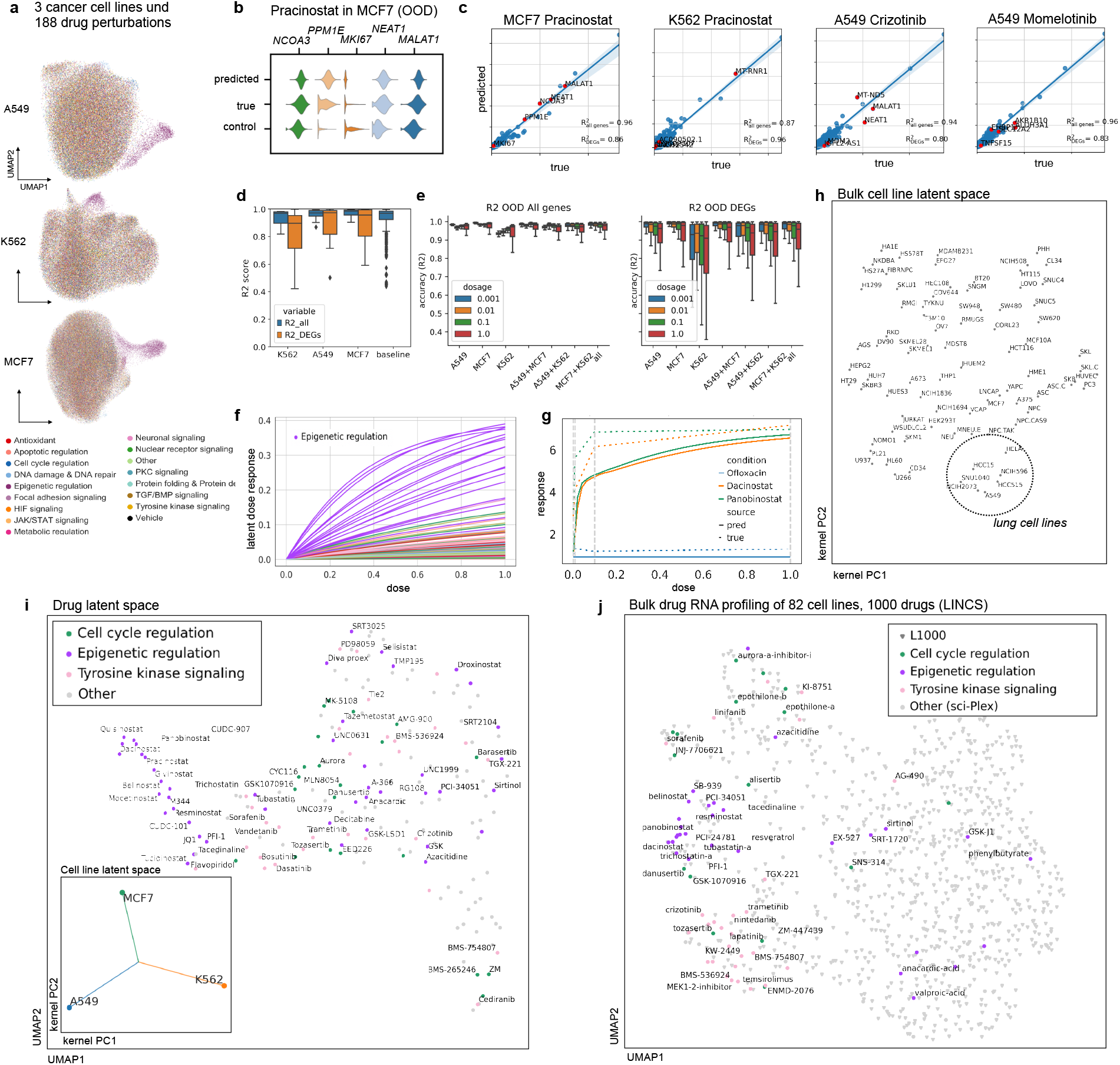
Learning drug and cell line latent representations from massive single-cell screens of 188 drugs across cancer cell lines. **(a)** UMAP representation of sci-Plex samples of A549, K562 and MCF7 cell-lines colored by pathway targeted by the compounds to which cells were exposed. **(b)** Distribution of top 5 differentially expressed genes in MCF7 cells after treatment with Pracinostat at the highest dose for real, control and CPA predicted cells. **(c)** Mean gene expression of 5,000 genes and top 50 DEGs between CPA predicted and real cells together with the top five DEGs highlighted in red for four compounds for which the model did not see any examples of the highest dose. **(d)** Box plots of *R*^2^ scores for predicted and real cells for 36 compounds and 108 unique held out perturbations across different cell lines. Baseline indicates comparison of real compounds with each other. **(e)** *R*^2^ scores box plot for all and top 50 DEGs. Each column represents a scenario where cells exposed with specific dose for all compounds on a cell line or combinations of cell lines were held from training and later predicted. **(f)** Latent dose response obtained from dose encoder for all compounds colored by pathways. **(g)** Real and predicted dose response curves based on gene expression data, for a single compound with differential dose response across three cell lines. **(h)** Latent representation of 80 cell lines from L1000 dataset. **(i)** Two dimensional representation of latent drug embeddings as learned by the CPA. Compounds associated with epigenetic regulation, tyrosine kinase signaling, and cell cycle regulation pathways are colored by their respective known pathways. The lower left panel shows latent covariate embedding for three cell lines in the data, indicating no specific similarity preference. **(j)** Latent drug embedding of CPA model trained on the bulk-RNA cell line profiles from the L1000 dataset, with focus on drugs shared with the sci-Plex experiment from (a).

CPA is able to extrapolate to the unseen OOD conditions with unexpected accuracy, as it captures the difference between control and treated conditions also for a compound where it did not see examples with the highest dose. As one example, pracinostat has a strong differential response to treatment compared to control, as can be seen from the distributions of the top 5 differentially expressed genes (**Figure 3b**). Despite not seeing the effect of Pracinostat at the highest dose in any of the three cell lines, CPA correctly infers the mean and distribution of these genes (**Figure 3b**). CPA performs well in modeling unseen perturbations, as the correlation of real and predicted values across OOD conditions is overall better than the correlation between real values (**Figure 3c**). When looking at individual conditions (**Figure 3d**), CPA does well recapitulating genes with low and high mean expression in the OOD condition.

CPA has lower performance when predicting experiments with more unseen covariates. To assess the ability of the model to generalize to unseen conditions, we trained CPA on 28 splits with different held-out conditions, with one of the doses held out in anywhere between 1-3 cell lines (**Figure 3e**). We see here that K562 is the hardest cell line to generalize, when considering training on two cell lines to generalize to another. We also see that extrapolating to the highest dose is a harder task than interpolating intermediate doses, which is consistent with the difficulty of anticipating the experimental effect of a higher dose, versus fitting sigmoidal behavior to model intermediate doses. When examining the shape of the sigmoid per compound learned by the model (**Figure 3f**), we see that epigenetic compounds, which caused the greatest differential expression effects, have higher latent response curves, indicating that CPA learns a general, cell-line agnostic response strength measure for compounds. This learned sigmoid behavior can then be used in conjunction with the latent vectors to reconstruct the gene expression of treated cells over interpolated doses (**Figure 3g**).

After training, CPA learns a compressed representation of the 188 compounds, where each drug is represented by a single 256 dimensional vector (**Figure 3i**). To test whether the learned drug embeddings are meaningful, we asked if compounds with similar putative mechanisms of action are similar in latent space. This holds for a large set of major mechanisms: we find that epigenetic, tyrosine kinase signaling, and cell-cycle regulation compounds are clustered together by the model, which suggests the effectiveness of drugs with these mechanisms on these three cancer cell-lines which is in line with the findings in the original publication [4].

We additionally demonstrate that the model learns universal relationships between compounds which remain true across datasets and modalities. Using the same set of compounds tested in the sci-Plex dataset together with 853 other compounds (for a total of 1000 compounds), we trained CPA on L1000 bulk perturbation measurement data across 82 cell lines [36]. We observed that CPA works equally well on bulk RNA-seq data, and also that matched epigenetic and tyrosine kinase signaling compounds present in sci-Plex were close to each other in the latent representation, suggesting that the learned model similarities apply across datasets (**Figure 3j**). This holds also for the other learned embeddings: Applying the same similarity metric to the covariate embedding-here the 82 cell lines – we observed that the cell line embedding learned by the model also represents cell line similarity in response to perturbation, as cell lines from lung tissue were clustered together (**Figure 3h**).

### CPA allows modeling combinatorial genetic perturbation patterns

Combinatorial drug therapies are hypothesized to address the limited effectiveness of mono-therapies[37] and prevent drug resistance in cancer therapies[37–39]. However, the combined expression of a small number of genes often drives the complexity at the cellular level, leading to the emergence of new properties, behaviors, and diverse cell types [5]. To study such genetic interactions (GIs), recent perturbation scRNA-seq assays allow us to measure the gene expression response of a cell to the perturbation of genes alone or in combination[12, 13]. While experimental approaches are necessary to assess the effect of combination therapies, in practice, it becomes infeasible to experimentally explore all possible combinations without computational predictions.

To pursue this aim, we applied our CPA model to scRNA-seq data collected from Perturb-seq (single-cell RNA-sequencing pooled CRISPR screens) to assess how overexpression of single or combinatorial interactions of 105 genes (i.e., single gene x, single gene y, and pair x+y) affected the growth of K562 cells [5]. In total, this dataset contains 284 conditions measured across 108, 000 single-cells, where 131 are unique combination pairs (i.e., x+y) and the rest are single gene perturbations or control cells. We observed that the latent genetic interaction manifold placed GIs inducing known and similar gene programs close to each other (**Figure 4a**). For example, consider *CBL* (orange cluster in **Figure 4a**): the surrounding points, comprising its regulators (e.g., *UBASH3A/B*) and multisubstrate tyrosine phosphatases (e.g., *PTPN9/12*), have all been previously reported to induce erythroid markers [5]. Next, we sought to assess our ability to predict specific genetic interactions. We examined a synergistic interaction between *CBL* and *CNN1* in driving erythroid differentiation which has been previously validated [5]. We trained a CPA model with *CBL+CNN1* held out from the training data. Overexpression of either gene leads to the progression of cells from control to single perturbed and doubly perturbed cells (**Supplementary Fig.2a**) toward the erythroid gene program. Overexpression of both *CBL* and *CNN1* up-regulate known gene markers[5] such as hemoglobins (see *HBA1/2* and *HBG1/2* in **Figure 4b**). We observed that our model successfully predicted this synergistic interaction, recapitulating patterns similar to real data and inline with the original findings (**Figure 4c**). We further evaluated CPA to predict a previously reported[5] genetic epistatic interaction between *DUSP9* and *ETS1*, leading to domination of the *DUSP9* phenotype in doubly perturbed cells (**Figure 4c**).

**Figure 4:**
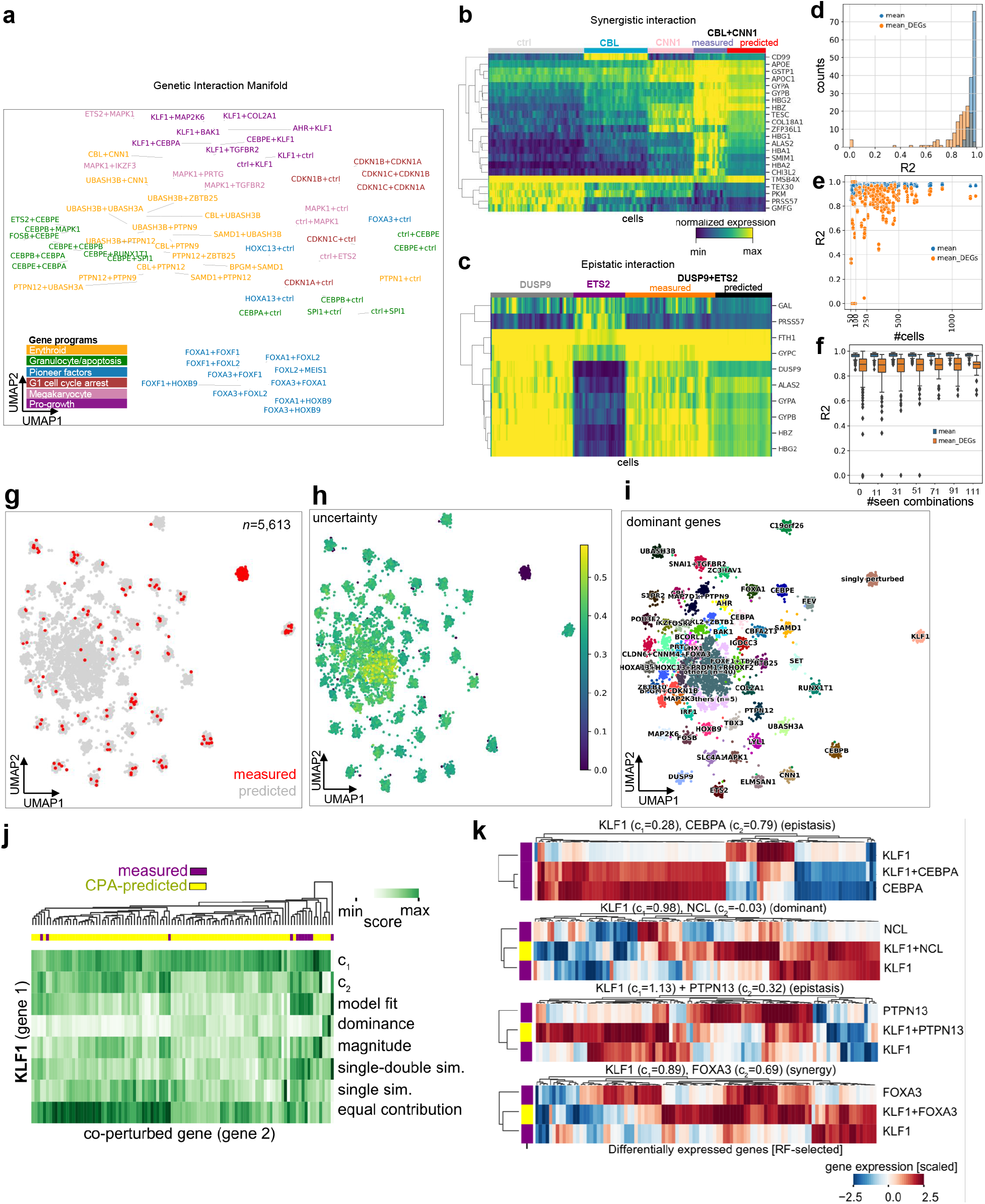
Learning and predicting combinatorial genetic perturbations. **(a)** UMAP inferred latent space using CPA for 281 single-and double-gene perturbations obtained from Perturb-seq[5]. Each dot represents a genetic perturbation. Coloring indicates gene programs associated to perturbed genes. **(b)** Measured and CPA-predicted gene expression for cells linked to a synergistic gene pair (*CBL+CNN1*). Gene names taken from the original publication. **(c)** As (b) for an epistatic (*DUSP9+ETS*) gene pair. Top 10 DEGs of *DUSP9+ETS* co-perturbed cells versus control cells are shown. **(d)** R2 values of mean gene-expression of measured and predicted cells for all genes (blue) or top 100 DEGs for the prediction of all 131 combinations (13 trained models, with 10 tested combinations each time) (orange). **(e)** R2 values of predicted and real mean gene-expression versus number of cells in the real data **(h)** R2 values for predicted and real cells versus number of combinations seen during training. **(g)** UMAP of measured (n=284, red dots) and CPA-predicted (n=5,329, gray dots) perturbation combinations. **(h)** As (g), showing measurement uncertainty (cosine similarity). **(i)** As (g), showing dominant genes in leiden clusters (25 or more observations).**(j)** Hierarchical clustering of linear regression associated metrics between *KLF1* with co-perturbed genes, in measured and predicted cells). **(k)** Scaled gene expression changes (versus control) of RF-selected genes (x-axis) in measured (purple) and predicted (yellow) perturbations (y-axis). Headers indicate gene-wise regression coefficients, and interaction mode suggestions[5].

To systemically evaluate the CPA’s generalization behavior, we trained 13 different models while leaving out all cells from ≈ 10 unique combinations covering all 131 doubly perturbed conditions in the dataset, which were predicted following training. The reported *R*^2^ values showed robust prediction for most of the perturbations: lower scores were seen for perturbations where the evaluation was noisy due to sample scarcity (n < 100), or when one of the perturbations was only available as singly perturbed cells in the data, leading the model to fail to predict the unseen combination (**Figure 4d-e**, see **Supplementary Fig. 2**). To further understand when CPA performance deteriorated, we first trained it on a subset with no combinations seen during training, and then gradually increased the number of combinations seen during training. We found that overall prediction accuracy improved when the model was trained with more combinations, and that it could fail to predict DEGs when trained with fewer combinations (see n < 71 combinations in **Figure 4f**).

Hence, once trained with sufficiently large and diverse training data, CPA could robustly predict unseen perturbations. We next asked whether our model could generalize beyond the measured combinations and generate *in-silico* all 5,329 combinations, which were not measured in the real dataset, but made up ≈ 98% of all possibilities. To study the quality of these predictions, we trained a model where all combinations were seen during training to achieve maximum training data and sample diversity. We then predicted 50 single-cells for all missing combinations. We found that, while the latent embeddings did not fully capture all the nuances in the similarity of perturbations compared to gene space, it provided an abstract and easier to perform high-level overview of potential perturbation combinations. Thus, we leveraged our latent space to co-embed (**Figure 4g**) all measured and generated data while proving an uncertainty metric based on the distance from the measured phenotypes (**Figure 4h**). We hypothesized that the closer the generated embedding was to the measured data. the more likely it was to explore a similar space of the genetic manifold around the measured data. Meanwhile, the distant points can potentially indicate novel behaviors, although this would require additional consideration and validation steps. Equipped with this information, we annotated the embedding clusters based on gene prevalence, finding that single genes (i.e. gene x) paired with other genes (i.e., y) as combinations (i.e., x+y) are a main driver of cluster separation (**Figure 4i**). Genes without measured double perturbations were less likely to be separated as independent clusters using the newly predicted transcriptomic expression (**Supplementary Fig. 3a**), suggesting that their interaction-specific effects were less variable than cases with at least one double perturbation available in the training data.

To investigate the type of interaction between the newly predicted conditions, we compared the differences between double and single perturbations versus control cells and thus annotated their interaction modes (adapted from [5] for *in silico* predictions). In each gene-specific cluster, we observed variability across these values, suggesting that our predictions contained granularity that went beyond single gene perturbation effects, and could not be fully dissected by two dimensional embeddings. Upon curation of gene perturbations using these metrics and the levels of experimental data available (**Supplementary Fig. 3b**), we decided to predict and annotate interaction modes based on these values when double measurements were available for at least one gene. For example, we observed clustering of *KLF1* and partner gene perturbation pairs solely from these metrics, suggesting the existence of several interaction modes (**Figure 4j**). When we further examine the differentially expressed genes in each co-perturbation, our metrics validated previously reported epistatic interactions (*CEBPA*), and proposed new ones with *KLF1*-dominant behavior (*NCL*), gene synergy (*FOXA3*), and epistasis (*PTPN13*), among others (**Figure 4k**). Repeating this analysis across all measured and predicted double perturbations, we found genes with potential interaction prevalences (**Supplementary Fig. 3c**). Among genes which repeatedly respond to several perturbations, we found common gene expression trends in both direction and magnitude (**Supplementary Fig. 3d**), suggesting that variation is modulated by conserved gene regulatory principles that are potentially captured in our learned model.

Altogether, our analysis indicated that double perturbation measurements can be generated by CPA by leveraging genetic perturbation data, which when combined with an uncertainty metric allows us to generate and interpret gene regulatory rules in the predicted gene-gene perturbations.

## Discussion

*In-silico* prediction of cell behavior in response to a perturbation is critical for optimal experiment design and the identification of effective drugs and treatments. With CPA, we have introduced a versatile and interpretable approach to modeling cell behaviors at single-cell resolution. CPA is implemented as a neural network trained using stochastic gradient descent, scaling up to millions of cells and thousands of genes.

We applied CPA to a variety of datasets and tasks, from predicting single-cell responses to learning embeddings, as well as reconstructing the expression response of compounds, with variable drug-dose combinations. Specifically, we illustrated the modeling of perturbations across dosage levels and time series, and have demonstrated applications in drug perturbation studies, as well as genetic perturbation assays with multiple gene knockouts, revealing potential gene-gene interaction modes inferred by our model predicted values. CPA combines the interpretability of linear decomposition models with the flexibility of nonlinear embedding models.

While CPA performed well in our experiments, it is well known that in machine learning there is no free lunch, and as with any other machine learning model, CPA will fail if the test data are very different from the training data. To alert CPA users to these cases, it is crucial to quantify model uncertainty. To do so, we implemented a distance-based uncertainty score to evaluate our predictions. Additionally, scalable Bayesian uncertainty models are promising alternatives for future work[40]. Although we opted to implement a deterministic autoencoder scheme, extensions towards variational models[17, 23], as well as cost functions other than mean squared error[22] are straightforward.

Aside from CPA, existing methods[17, 26] such as scGen[16] have also been shown capable of predicting single-cell perturbation responses when the dataset contains no combinatorial treatment or dose-dependent perturbations. Therefore, it may be beneficial to benchmark CPA against such methods on less complicated scenarios with few perturbations. However, this approach might not be practical, considering the current trend towards the generation of massive perturbation studies[4, 5, 12].

Currently, the model is based on gene expression alone, so it cannot directly capture other levels of interactions or effects, such as those due to post-transcriptional modification, signaling, or cell communication. However, due to the flexibility of neural network-based approaches, CPA could be extended to include other modalities, for example via multimodal single-cell CRISPR[41, 42] combined scRNA-and ATAC-seq[43, 44] and CUT&Tag[45, 46]. In particular, we expect spatial transcriptomics[47, 48] to be a valuable source for extensions to CPA due to its high sample number and the dominance of DL models in computer vision.

The CPA model is not limited to single-cell perturbations. While we chose the single-cell setting due to the high sample numbers available, the CPA could readily be applied to large-scale bulk cohorts, in which covariates might be patient ID or transcription factor perturbation. These and any other available attributes could be controlled independently[31] to achieve compositional, interpretable predictions. Any bulk compositional model may be combined with a smaller-scale single-cell model to compose truly multi-scale models of observed variance. The flexibility of the DL setting will also allow addition of constraints on perturbation or covariate latent spaces. These could, for example, be the similarity of chemical compounds[49], or clinical-covariate induced differences of patient IDs. The key feature of the CPA versus a normal autoencoder is its latent space disentanglement and the induced interpretability of the perturbations in the context of cell states and covariates. Eventually, any aim in biology is not only blind prediction, but mechanistic understanding. This objective is exemplified by the direction that DL models are taking in sequence genomics, where the aim is not only the prediction of new interactions, but also the interpretation of the learned gene regulation code. We therefore believe that CPA can not only be used as a hypothesis generation tool for *in-silico* screens but also as an overall data approximation model. Deviations from our assumed data generation process (see **Methods**) would then tell us about missing information in the given data set and/or missing aspects in the factor model. By including multiple layers of regulation, the resulting model can grow in flexibility for prediction and for mechanistic understanding on for example synergistic gene regulation or other interactions.

Finally, we expect CPA to facilitate new opportunities in expression-based perturbation screening, not only to learn optimal drug combinations, but also in how to personalize experiments and treatments by tailoring them based on individual cell response.

## Code availability

Code to reproduce all of our results is available at http://github.com/facebookresearch/CPA.

## Data availability

All datasets analyzed in this manuscript are public and have published in other papers. We have referenced them in the manuscript and made available at http://github.com/facebookresearch/CPA.

## Author Contributions

M.L., A.K.S, and D.L.P. conceived the project with contributions from F.J.T. D.L.P., M.L. and A.K.S designed the algorithm and implemented the first version. Y.J., C.D. and A.K.S. performed the first refactor. The final code is implemented by D.L.P. and A.K.S. with contributions from C.D., Y.J. and M.L. M.L. and C.D. curated all the datasets. F.A.W. helped interpret the model and results. M.L., A.K.S. and F.J.T. designed analyses and use-cases. M.L., A.K.S., Y.J., I.L.I. and C.D performed the analysis. F.J.T. and N.Y. supervised the research. All authors wrote the manuscript.

## Acknowledgments

M.L. and F.J.T. are grateful for valuable feedback from Aviv Regev and Dana Pe’er. We appreciate support from all members of Theis lab, specifically Malte D. Luecken and Fabiola Curion for their feedback and proof-reading. M.L is thankful for early graphical designs by Monir Jazaeri (Jaz) which did not make it to the final version of the paper. F.J.T. acknowledges support by the BMBF (grant L031L0214A, grant 01IS18036A and grant 01IS18053A), by the Helmholtz Association (Incubator grant sparse2big, grant ZT-I-0007) and by the Chan Zuckerberg Initiative DAF (advised fund of Silicon Valley Community Foundation, 2018-182835 and 2019-207271). This work was further supported by Helmholtz Association’s Initiative and Networking Fund through Helmholtz AI [grant ZT-I-PF-5-01]. I.L.I. has received funding from the European Union’s Horizon 2020 research and innovation programme under grant agreement No 874656.

## Competing interests

F.J.T. reports receiving consulting fees from Roche Diagnostics GmbH and Cellarity Inc., and ownership interest in Cellarity, Inc. and Dermagnostix. F.A.W. is a full-time employee of Cellarity Inc., and has ownership interest in Cellarity, Inc.

## Methods

### Data generating process

We consider a dataset 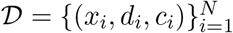 where each *x_i_* ∈ ℝ^*G*^ describes the gene expression of *G* genes from cell *i*. The perturbation vector *d_i_* = (*d*_*i*,1_,…,*d*_*i,M*_) contains elements *d_i,j_* ≥0 describing the dose of drug *j* applied to cell *i*. If *d_i,j_* = 0, this means that perturbation *j* was not applied to cell *i*. Unless stated otherwise, the sequel assumes column vectors. Similarly, the vector of vectors *c_i_* = (*c*_*i*,1_,…*c_i,K_*) contains additional discrete covariates such as cell-types or species, where each covariate is itself a vector. More specifically, *c_i,j_* is a *K_j_*-dimensional one-hot vector.

We assume that an unknown generative model produced our data set 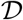. The three initial components of this generative process are a latent (unobserved) basal latent state 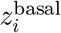 for cell *i*, together with its perturbation vector *d_i_* and covariate vector *c_i_*. We assume that the basal latent state is independent from the perturbation vector *d_i_* and covariate vector *c_i_*. Next, we form the latent (also unobserved) perturbed latent state *z_i_* as:

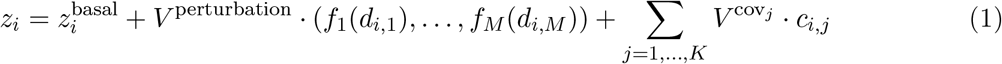

In this equation, each column of the matrix *V*^perturbation^ ∈ ℝ^*d×M*^ represents a *d*-dimensional embedding for one of the *M* possible perturbations represented in *d_i_*. Similarly, each column of the matrix *V*^cov_*j*_^ ∈ ℝ^*d×K_j_*^ represents a *d*-dimensional embedding for the *j*-th discrete covariate, represented as a *K_j_*-dimensional one-hot vector *c_i,j_*. The functions *f_j_*: ℝ → ℝ scale non-linearly each of the *d_i,j_* in the perturbation vector *d_i_*, therefore implementing *M* independent dose-response (or time-response) curves. Finally, we assume that the generative process returns the observed gene expression *x_i_* by means of an unknown decoding distribution *p*(*x_i_*|*z_i_*). This process builds the observation (*x_i_, d_i_, c_i_*), which is then included in our dataset 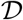.

### Compositional Perturbation Autoencoder (CPA)

Assuming the generative process described above, our goal is to train a machine learning model 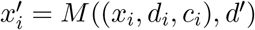 such that, given a dataset triplet (*x_i_, d_i_, c_i_*) as well as a target perturbation *d*‛ estimates the gene expression 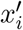. The term 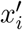 represents what would the counterfactual distribution of the gene expression *x_i_* with covariates *c_i_* look like, had it been perturbed with *d*‛ instead of *d_i_*.

Given a dataset and a learning goal, we are now ready to describe our proposed model, the Compositional Perturbation Autoencoder (CPA). In the following, we describe separately how to train and test CPA models.

### Training

The training of a CPA model consists in auto-encoding dataset triplets (*x_i_, d_i_, c_i_*). That is, during training, a CPA model does not attempt to answer counterfactual questions. Instead, the training process consists in (1) encoding the gene expression *x_i_* into an estimated basal state 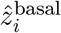 that does not contain any information about (*d_i_, c_i_*), (2) combining 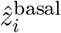 with learnable embeddings about (*d_i_, c_i_*) to form an estimated perturbed state 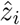 and (3) decoding 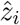 back into the observed gene expression *x_i_*.

More specifically, the CPA model first encodes the observed gene expression *x_i_* into an estimated basal state:

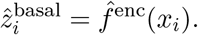

In turn, the estimated basal state is used to compute the estimated perturbed state 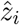:

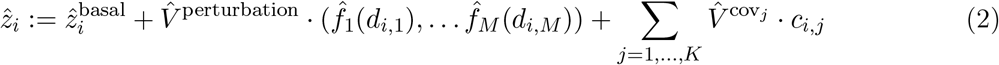

Contrary to (1), this expression introduces three additional learnable components: the perturbation embeddings 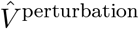, the covariate embeddings 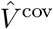 and the learnable dose-response curves 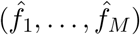, here implemented as small neural networks constrained to satisfy 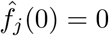.

As a final step, a decoder 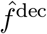 accepts the estimated perturbed state 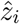 and returns 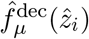 and 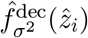, that is, the estimated mean and variance of the counterfactual gene expression 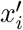.

To train CPA models, we use three loss functions. First, the reconstruction loss function is the Gaussian negative log-likelihood:

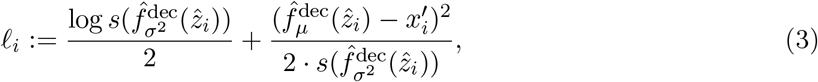
where *s*(*σ*^2^) = log(exp(*σ*^2^ + 10^−3^) + 1) enforces a positivity constraint on the variance and adds numerical stability. This loss function rewards the end-to-end auto-encoding process if producing the observed gene expression *x_i_*.

Second, and according to our assumptions about the data generating process, we are interested in removing the information about (*d_i_*; *c_i_*) from 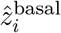. To achieve this information removal, we follow an adversarial approach [31]. In particular, we set up the following auxiliary loss functions:

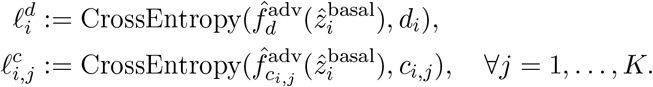

The functions 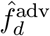, 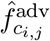 are a collection of neural network classifiers trying to predict about (*d_i_*; *c_i_*) given the estimated basal state 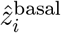.

Given this collection of losses, the training process is an optimization problem that repeats the following two steps:

1. sample 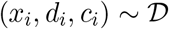, minimize 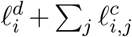 by updating the parameters of 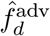 and 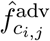 for all *j* = 1,…,*K*;
2. sample 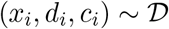, minimize 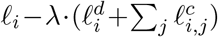 by updating the parameters of the encoder 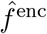, the decoder 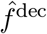, the perturbation embeddings 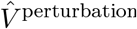 the covariate embeddings 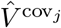 for all *j* = 1,…,*K*, and the dose-response curve estimators 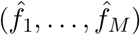

### Testing

Given an observation (*x_i_*; *d_i_*; *c_i_*) and a counterfactual treatment *d*‛, we can use a trained CPA model to answer what would the counterfactual distribution of the gene expression *x_i_* with covariates *c_i_* look like, had it been perturbed with *d*‛ instead of *d_i_*. To this end, we follow the following process:

1. Compute the estimated basal state:
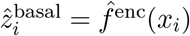
2. Compute the counterfactual perturbed state 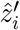

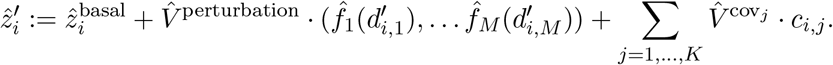 Note that in the previous expression, we are using the counterfactual treatment *d*‛ instead of the observed treatment *d_i_*.
3. Compute and return the counterfactual gene expression mean 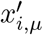:

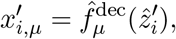
and variance 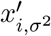:

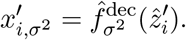

### Hyper-parameters and training

For each dataset, we perform a random hyper-parameter search of 100 trials. The table below outlines the distribution of values for each of the hyper-parameters involved in CPA training.

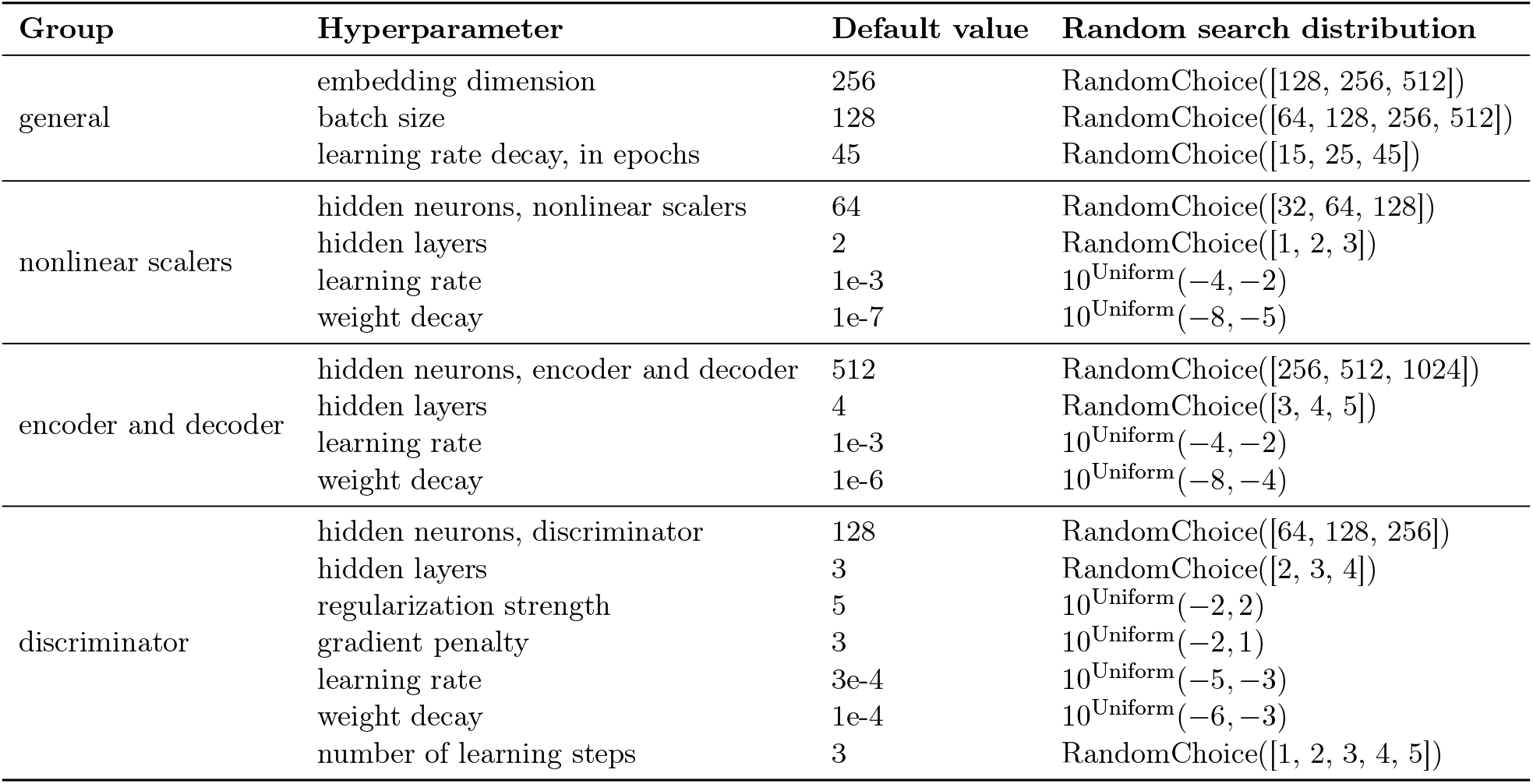

### Model evaluation

We use several metrics to evaluate the performance of our model: (1) quality of reconstruction for in and OOD cases and (2) quality of disentanglement of cell information from perturbation information.

We split each dataset into 3 subsets: train, test, and OOD. For OOD cases, we choose combinations of perturbations that exhibit unseen behavior. This usually corresponds to the most extreme drug dosages. We select one perturbation combination as “control”. Usually these are Vehicle or DMSO if real control samples are present in the dataset, otherwise we choose a drug perturbation at a lower dosage as “control”. For the evaluation, we use the mean squared error of the reconstruction of an individual cell and average it over the cells for the perturbation of interest. As an additional metric we use classification accuracy in order to check how well the information about the drugs was separated from the information about the cells.

### Uncertainty estimation

To estimate the uncertainty of the predictions we use as a proxy the minimum distance between the queried perturbation and the set of conditions (covariate + perturbation combinations) seen during training (Supplementary Fig.1). Intuitively, we expect predictions on queried conditions that are more distant from the set of seen conditions to be more uncertain. To estimate this distance we first compute the set of embeddings of the training covariate and perturbation combinations:

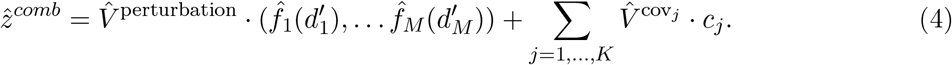

The latent vector for the queried condition is obtained in the same manner. The cosine and euclidean distances from the training embedding set are computed and the minimum distance is used as a proxy for uncertainty.

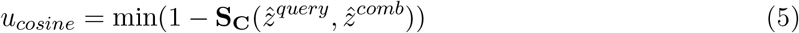

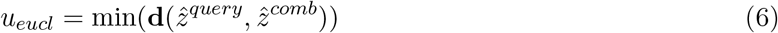

Where S_C_(*x*; *y*) stands for the cosine similarity and d(*x*; *y*) for the euclidean distance between the two vectors.

With this methodology, in the case of a drug screening experiment, if we query a combination of cell type, drug, and dosage that was seen during training, we get an uncertainty of zero, since this combination was present in the training set. It is important to note that with this method we obtain a condition-level uncertainty, in that all cells predicted under the same query will have the same uncertainty, thus not taking cell specific information into account.

### R2 score

We used the *r^2^_score* function from *scikit-learn* which reports R2 (coefficient of determination) regression score.

### Datasets

#### Gehring et al

This dataset[8] comprises of 21; 191 neural stem cells (NSCs) cells perturbed with EGF/bFGF, BMP4, decitabine, scriptaid, and retinoic acid. We obtained normalized data from the original authors and after QC filtering 19; 637 cells remained. We further selected 5; 000 highly variable genes (HVGs) using SCANPY’s[50] *highly_variable_genes* function for training and evaluation of the model.

#### Genetic CRISPR screening experiment

We obtained the raw count matrices from Norman et al.[5] from GEO (accession ID GSE133344). According to authors guide, we excluded “NegCtrl1_NegCtrl0__NegCtrl1_NegCtrl0” control cells and merged all unperturbed cells as one “ctrl” condition. We then normalized and log-transformed the data using SCANPY and selected 5; 000 HVGs for training. The processed dataset contained 108; 497 cells.

#### Cross-species experiment

The data was generated by Hagai et al.[15] and downloaded from ArrayExpress (accession: E-MTAB-6754). The data consists of 119; 819 phagocytes obtained from four different species: mouse, rat, pig and rabbit. Phagocytes were treated with lipopolysaccharide (LPS) and the samples were collected at different time points: 0 (control), 2, 4, and 6 hours after the beginning of treatment. All genes from non-mouse data were mapped to the respective orthologs in the mouse genome using Ensembl ID annotations. We filtered out cells with a percentage of counts belonging to mitochondrial genes higher than 20%, then proceeded to normalize and log-transform the count data. For training and evaluation, we selected 5000 HVG using SCANPY. After filtering, the data consists of 113; 400 cells.

#### sci-Plex 2

The data was generated by Srivatsan et al. [35] and downloaded from GEO (GSM4150377). The dataset consists of A549 cells treated with one of the following four compounds: dexamethasone, Nutlin-3a, BMS-345541, or vorinostat (SAHA). The treatment lasted 24 hours across seven different doses. The count matrix obtained from GEO consists of 24; 262 cells. During QC we filtered cells with fewer than 500 counts and 720 detected genes. We discarded cells with a percentage of mitochondrial gene counts higher than 10%, thus reducing the dataset to 14; 811 cells. Genes present in fewer than 100 cells were discarded. We normalized the data using the size factors provided by the authors and log-transformed it. We selected 5000 HVGs for training and further evaluations.

#### sci-Plex 3

The data was generated by Srivatsan et al.[35] and downloaded from GEO (GSM4150378). The dataset consists of three cancer cell lines (A549, MCF7, K562), which are treated with 188 different compounds with different mechanisms of action. The cells are treated with 4 dosages (10, 100, 1000, and 10000 nM) plus vehicle. The count matrix obtained from GEO consists of 581,777 cells. The data was subset to half its size, reducing it to 290,888 cells. We then proceeded with log-transformation and the the selection of 5000 HVGs using SCANPY.

### Interpretation of combinatorial genetic interactions by perturbation pairs and responder genes

In the case of genetic screening, previous work by [5] proposed a set of metrics to annotate and classify gene-gene interactions based on responder genes. Based on this, here we used measured or predicted gene expression differences with respect to control cells (*δ*), for gene perturbations a (*δa*), b (*δb*) and double perturbations ab (*δab*), to calculate interaction types by similarity between these three expression vectors.

More specifically, to calculate association coefficients, we use the linear regression coefficients *c*_1_ and *c*_2_ obtained from the model

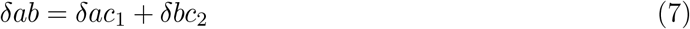

To describe interaction modes, we used the following metrics.

1. **similarity between predicted and observed values**: dcor(*δac*_1_ + *δbc*_2_, *δab*).
2. **linear regression coefficients**: *c*_1_ and *c*_2_.
3. **magnitude**: 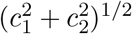
4. **dominance**: |*log*_10_(*c*_1_=*c*_2_)|.
5. **similarity of single transcriptomes**: *dcor*(*a*; *b*)
6. **similarity of single to double transcriptomes**: *dcor*([*a*; *b*]; *ab*).
7. **equal contributions**: 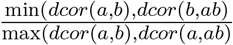

Following clustering and comparison of these metrics across measured and predicted cells, we decided the following rules of thumb to define and annotate interaction modes:

1. **Epistatic:** min(*abs*(*c*_1_); *abs*(*c*_2_)) > 0:2 and either (i) (*abs*(*c*_1_) > 2*abs*(*c*_2_)) or (ii) (*abs*(*c*_2_) > 2abs(*c*_1_))
2. **potentiation:** magnitude > 1 and abs(dcor(*a*; *b*))-1 > 0.2.
3. **strong sinergy (similar phenotypes):** magnitude > 1 and abs(*dcor*([*a*, *b*]; *ab*))-1 > 0.2
4. **strong sinergy (different phenotypes):** magnitude > 1 and abs(*dcor*(*a*, *b*))-1 > 0.5.
5. **additive:** abs(magnitude)-1 < 0.1.
6. **redundant:** abs(*dcor*([*a*; *b*]; *ab*))-1 < 0.2 and abs(*dcor*(*a*, *b*))-1 < 0.2

More than one genetic interaction can be suggested from these rules. In those cases, we did not assign any plausible interaction. For visualization purposes, we consider perturbed genes with 50 or more interaction modes reported with other co-perturbed genes (Supplementary Fig.3c).

To visualize differentially expressed genes with similar response across perturbations (Supplementary Fig.3d), we trained a random forest classifier using as prediction labels control, a, b and ab cells, and gene expression as features. We retrieved the top 200 genes from this approach. Then, we annotated the direction (positive or negative) and the magnitude of those changes versus control cells, generating a code for clustering and visualization. To label genes with potential interaction effects, we labeled them if the double perturbation predicted magnitude is 1.5x times or higher than the best value observed in single perturbations.

---

**Supplementary Figure 1:**
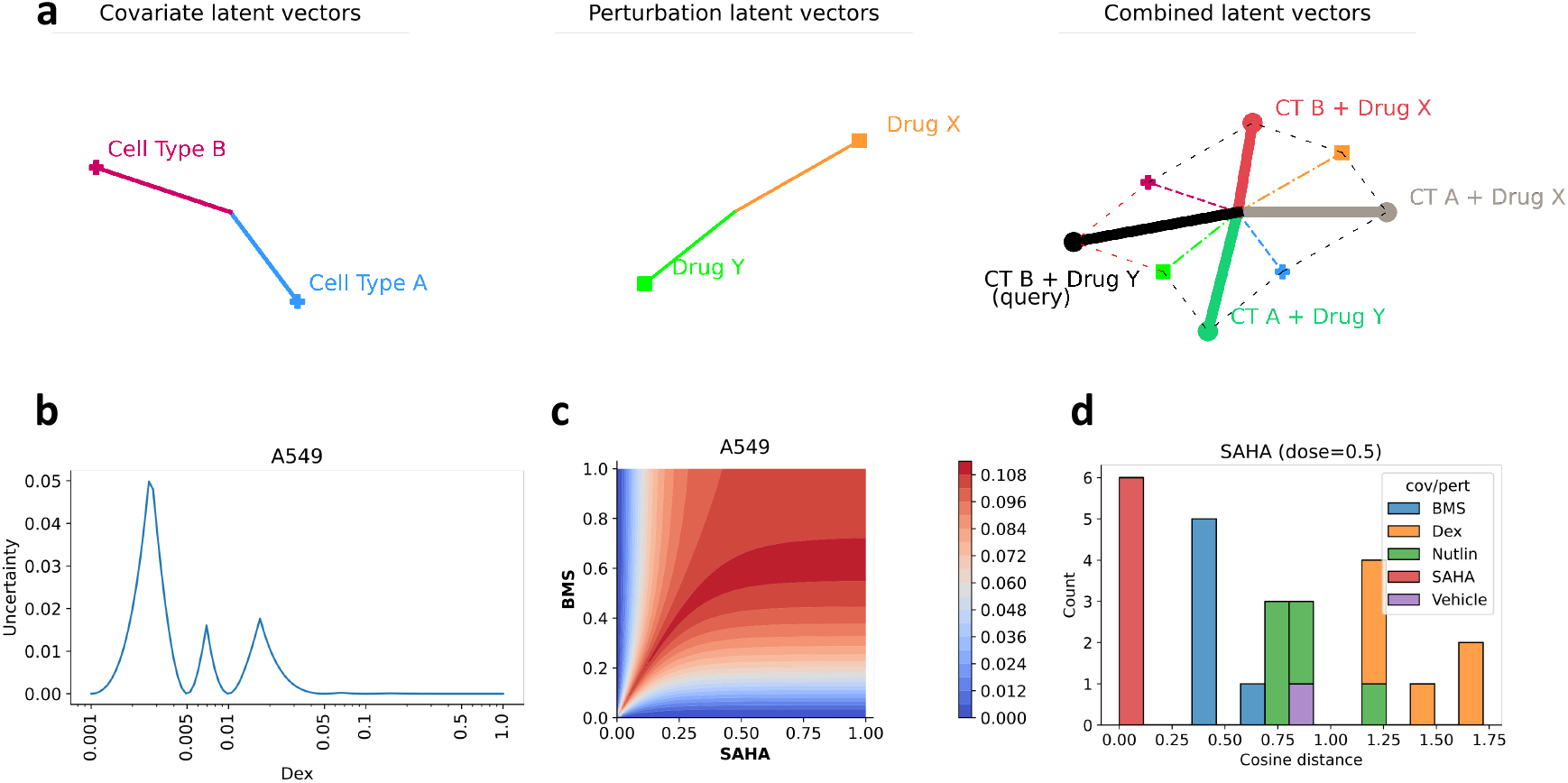
CPA uncertainty estimation. **(a)** Schematic representation of the steps involved in uncertainty estimation in the case of a dataset with two cell types and two drugs (single dosage per drug). The covariate and perturbation latent vectors are summed in order to generate the set of combinations in the training set. The distances from the query vector and all the vectors in the set are then computed. The closest distance is used as a proxy for uncertainty in the prediction of the model. **(b)** Example of uncertainty across dosages of Dexamethasone in the sci-Plex 2 dataset. The ticks on the x-axis (log-scaled) indicate dosages seen at training time for which the uncertainty is 0. The dosages were min-max normalized. **(c)** 2D plot of uncertainty across dosages (min-max normalized) of two different drugs and combinations thereof in the sci-Plex 2 dataset. **(d)** Example histogram of cosine distances between the SAHA (dose=0.5) and the vectors in the set of training perturbations. The distribution shows that training vectors belonging to the same perturbation but with different dosages have the lowest uncertainties, with other drugs being increasingly more distant.

**Supplementary Figure 2:**
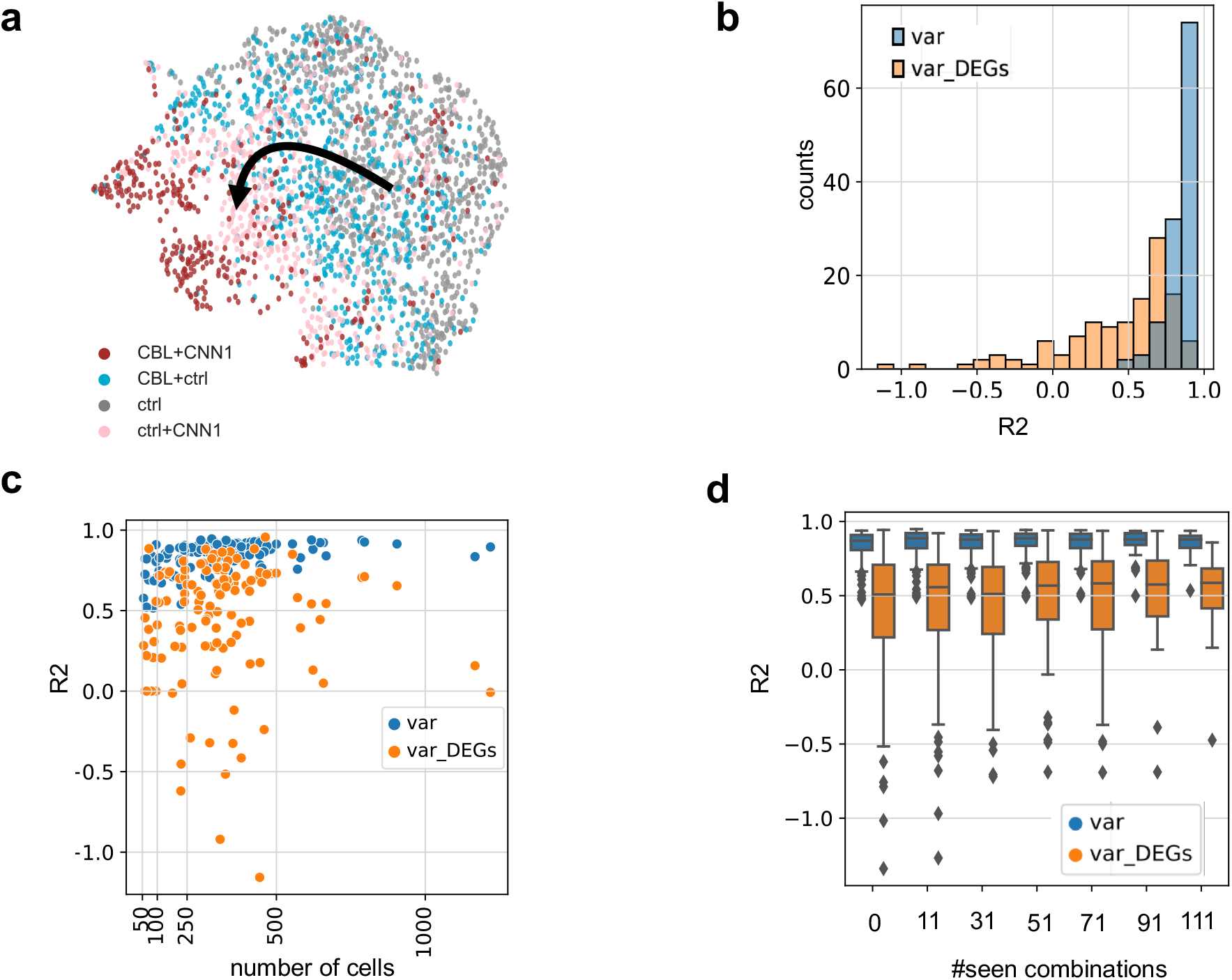
Performance evaluation for CPA combinatorial predictions. **(a)** UMAP representation of control (ctrl), singly perturbed (CBL+ctrl, ctrl+CNN1) and doubly (CBL+CNN1) perturbed cells. **(b)** *R*^2^ scores for all genes (blue) or top 100 DEGs (orange) for the prediction of all 131 combinations in the data by training 13 different models and leaving out ≈ 10 combinations each time. **(c)** Scatter plots of number of samples in the real data for each combination (x-axis) versus *R*^2^ values for the variance of predicted and real for that combination **(d)** Box-plots of *R*^2^ values for variance for predicted and real cells while increasing the number of combinations seen during training.

**Supplementary Figure 3:**
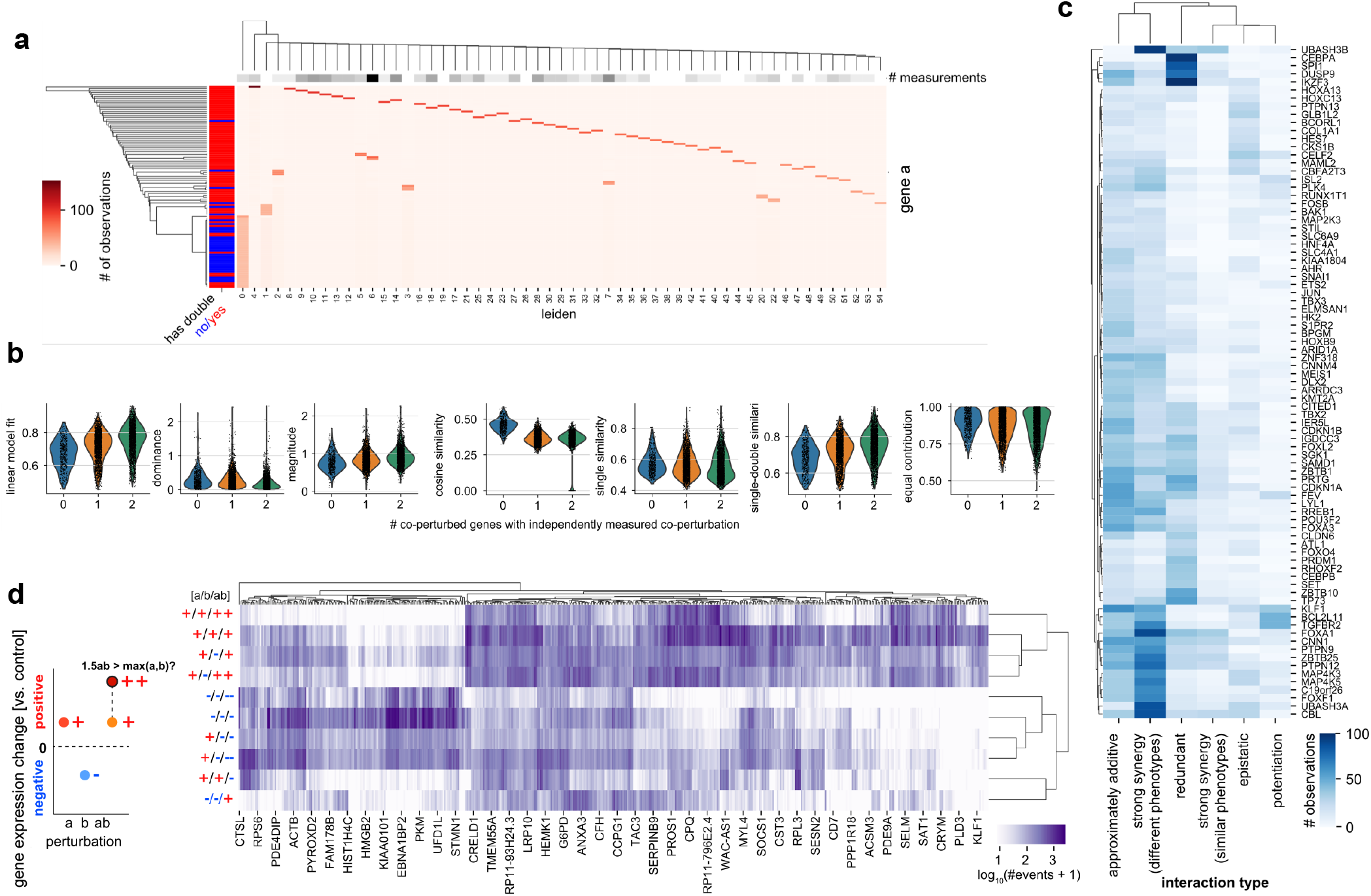
Gene-gene interaction insights revealed from genetic perturbation predictions using CPA. **(a)** Number of single gene observations in leiden clusters for generated measurements (from Figure 4i). Most leiden clusters contain a prevalence for one perturbed gene. The majority of genes without measured double perturbations share a limited number of clusters. **(b)** Quality control and interaction metrics to compare gene expression differences between single and double perturbations. Metrics vary based on number of genes with a measured double perturbation (See Methods for definitions). **(c)** Interaction mode counts predicted for all genes based on interaction metrics (based on [5]). **(d)** (*left*) Gene expression changes for double perturbations (ab) versus single perturbations (a, b), are compared by direction and magnitude. Positive (+) and negative (-) labels indicate increase/decrease versus control cells, and double positive/negative (++/−−) indicate values higher than 1.5 times the highest comparable value in single perturbations. (*right*) 500 genes with highest prevalence in differentially expressed genes across datasets, clustered by prevalent response types from single and double perturbations.

**Supplementary Table 1.** Performance scores for the sci-Plex 2 dataset. To improve readability the columns are sorted by: condition (first priority) and scores (second priority).

| condition | dose_val | R2_mean | R2_mean_DE | R2_var | split | num_cells |
| --- | --- | --- | --- | --- | --- | --- |
| SAHA | 0.01 | 0.99 | 0.99 | 0.95 | test | 160 |
| SAHA | 0.005 | 0.98 | 0.98 | 0.93 | test | 143 |
| SAHA | 0.05 | 0.98 | 0.95 | 0.93 | test | 118 |
| SAHA | 1.0 | 0.98 | 0.95 | 0.91 | test | 137 |
| SAHA | 0.001 | 0.97 | 0.97 | 0.92 | test | 169 |
| SAHA | 0.1 | 0.96 | 0.86 | 0.94 | test | 129 |
| SAHA | 0.5 | 0.96 | 0.86 | 0.89 | **ood** | 604 |
| Nutlin | 0.001 | 0.98 | 0.98 | 0.94 | test | 135 |
| Nutlin | 0.05 | 0.98 | 0.98 | 0.94 | test | 136 |
| Nutlin | 0.005 | 0.98 | 0.97 | 0.94 | test | 107 |
| Nutlin | 0.1 | 0.98 | 0.97 | 0.94 | test | 200 |
| Nutlin | 0.01 | 0.98 | 0.97 | 0.93 | test | 180 |
| Nutlin | 0.5 | 0.92 | 0.86 | 0.84 | **ood** | 265 |
| Nutlin | 1.0 | 0.26 | 0.61 | 0.00 | test | 1 |
| Dex | 0.5 | 0.99 | 0.99 | 0.98 | **ood** | 864 |
| Dex | 1.0 | 0.99 | 0.98 | 0.95 | test | 222 |
| Dex | 0.1 | 0.99 | 0.94 | 0.96 | test | 218 |
| Dex | 0.05 | 0.98 | 0.90 | 0.93 | test | 210 |
| Dex | 0.001 | 0.95 | 0.87 | 0.89 | test | 123 |
| Dex | 0.01 | 0.95 | 0.61 | 0.89 | test | 238 |
| Dex | 0.005 | 0.94 | 0.53 | 0.88 | test | 108 |
| BMS | 0.001 | 0.98 | 0.97 | 0.92 | test | 212 |
| BMS | 0.005 | 0.97 | 0.99 | 0.92 | test | 151 |
| BMS | 0.5 | 0.95 | 0.89 | 0.78 | **ood** | 34 |
| BMS | 0.01 | 0.95 | 0.80 | 0.86 | test | 82 |
| BMS | 0.05 | 0.93 | 0.87 | 0.75 | test | 59 |
| BMS | 0.1 | 0.92 | 0.89 | 0.74 | test | 50 |
| BMS | 1.0 | 0.55 | -0.87 | 0.21 | test | 6 |

**Supplementary Table 2.** A simple benchmark on OOD split for the sci-Plex 2 dataset.

| condition | R2_mean | R2_mean_DE | method |
| --- | --- | --- | --- |
| SAHA | 0.98 | 0.94 | linear |
| SAHA | 0.96 | 0.86 | CPA |
| Nutlin | 0.92 | 0.86 | CPA |
| Nutlin | 0.85 | 0.80 | linear |
| Dex | 1.00 | 1.00 | linear |
| Dex | 0.99 | 0.99 | CPA |
| BMS | 0.95 | 0.89 | CPA |
| BMS | 0.89 | 0.85 | linear |

**Supplementary Table 3.** Performance scores for the 96-plex-scRNAseq dataset. For the readability the columns are sorted by: condition (first priority) and scores (second priority).

| condition | dose_val | R2_mean | R2_mean_DE | R2_var | split | num_cells |
| --- | --- | --- | --- | --- | --- | --- |
| EGF+RA | 0.2+1.0 | 0.98 | 0.95 | 0.84 | test | 553 |
| EGF+RA | 1.0+1.0 | 0.97 | 0.94 | 0.67 | test | 199 |
| EGF+RA | 0.2+0.2 | 0.96 | 0.98 | 0.89 | test | 87 |
| EGF+RA | 0.04+1.0 | 0.96 | 0.90 | 0.60 | test | 54 |
| EGF+RA | 1.0+0.2 | 0.95 | 0.91 | 0.75 | test | 30 |
| EGF | 0.2 | 0.98 | 0.96 | 0.86 | test | 90 |
| EGF | 1.0 | 0.94 | 0.71 | 0.60 | test | 73 |
| BMP+RA | 0.2+1.0 | 0.91 | 0.84 | 0.60 | test | 22 |
| BMP+EGF+ScripDec | 0.2+0.2+1.0 | 0.97 | 0.97 | 0.82 | **ood** | 166 |
| BMP+EGF+ScripDec | 0.04+0.04+1.0 | 0.97 | 0.96 | 0.61 | test | 28 |
| BMP+EGF+ScripDec | 0.04+0.2+1.0 | 0.97 | 0.94 | 0.50 | test | 39 |
| BMP+EGF+ScripDec | 0.2+0.2+0.2 | 0.97 | 0.92 | 0.65 | **ood** | 304 |
| BMP+EGF+ScripDec | 0.04+0.2+0.2 | 0.97 | 0.91 | 0.74 | test | 33 |
| BMP+EGF+ScripDec | 0.2+0.04+1.0 | 0.96 | 0.96 | 0.26 | test | 32 |
| BMP+EGF+ScripDec | 0.2+0.04+0.2 | 0.96 | 0.94 | 0.34 | test | 20 |
| BMP+EGF+ScripDec | 1.0+0.2+0.2 | 0.96 | 0.91 | 0.74 | **ood** | 113 |
| BMP+EGF+ScripDec | 0.2+1.0+1.0 | 0.95 | 0.89 | 0.51 | **ood** | 112 |
| BMP+EGF+ScripDec | 0.04+0.04+0.2 | 0.95 | 0.87 | 0.52 | test | 19 |
| BMP+EGF+ScripDec | 0.2+1.0+0.2 | 0.95 | 0.83 | 0.58 | **ood** | 105 |
| BMP+EGF+ScripDec | 0.04+1.0+1.0 | 0.94 | 0.96 | 0.57 | test | 17 |
| BMP+EGF+ScripDec | 1.0+0.04+0.2 | 0.91 | 0.86 | 0.51 | test | 15 |
| BMP+EGF+RA | 0.04+1.0+0.2 | 0.98 | 0.99 | 0.85 | test | 50 |
| BMP+EGF+RA | 0.2+0.04+0.2 | 0.98 | 0.98 | 0.74 | test | 63 |
| BMP+EGF+RA | 0.04+1.0+1.0 | 0.98 | 0.97 | 0.86 | test | 198 |
| BMP+EGF+RA | 0.04+0.2+1.0 | 0.97 | 0.96 | 0.88 | test | 552 |
| BMP+EGF+RA | 0.2+0.04+1.0 | 0.97 | 0.96 | 0.70 | test | 48 |
| BMP+EGF+RA | 0.04+0.2+0.2 | 0.97 | 0.92 | 0.80 | test | 73 |
| BMP+EGF+RA | 0.2+0.2+0.2 | 0.97 | 0.88 | 0.52 | **ood** | 206 |
| BMP+EGF+RA | 0.04+0.04+0.2 | 0.96 | 0.96 | 0.73 | test | 24 |
| BMP+EGF+RA | 0.2+1.0+0.2 | 0.96 | 0.89 | 0.66 | **ood** | 216 |
| BMP+EGF+RA | 0.2+1.0+1.0 | 0.96 | 0.87 | 0.66 | **ood** | 147 |
| BMP+EGF+RA | 0.2+0.2+1.0 | 0.95 | 0.77 | 0.59 | **ood** | 132 |
| BMP+EGF | 0.04+0.2 | 0.97 | 0.98 | 0.78 | test | 96 |
| BMP+EGF | 0.04+1.0 | 0.97 | 0.92 | 0.81 | test | 209 |
| BMP+EGF | 0.04+0.04 | 0.97 | 0.92 | 0.75 | test | 39 |
| BMP+EGF | 1.0+0.04 | 0.95 | 0.86 | 0.33 | test | 19 |
| BMP+EGF | 1.0+1.0 | 0.94 | 0.75 | 0.06 | **ood** | 113 |

**Supplementary Table 4.** Performance scores for the cross-species dataset across different splits.

| split | species | time | R2_mean | R2_mean_DE | R2_var | num_cells |
| --- | --- | --- | --- | --- | --- | --- |
| split4 | rat | 6.0 | 0.97 | 0.91 | 0.85 | 7827 |
| split4 | rat | 6.0 | 0.96 | 0.86 | 0.89 | 7827 |
| split4 | mouse | 6.0 | 0.97 | 0.90 | 0.81 | 5280 |
| split4 | mouse | 6.0 | 0.96 | 0.85 | 0.93 | 5280 |
| split3 | rat | 6.0 | 0.89 | 0.70 | 0.72 | 7827 |
| split3 | rat | 6.0 | 0.86 | 0.55 | 0.40 | 7827 |
| split3 | rat | 4.0 | 0.95 | 0.80 | 0.80 | 5755 |
| split3 | rat | 4.0 | 0.94 | 0.77 | 0.63 | 5755 |
| split2 | rat | 6.0 | 0.55 | -0.65 | -2.18 | 7827 |
| split2 | rat | 6.0 | 0.54 | 0.02 | 0.02 | 7827 |
| split2 | rat | 4.0 | 0.74 | -0.47 | 0.26 | 5755 |
| split2 | rat | 4.0 | 0.39 | -0.85 | -0.47 | 5755 |
| split2 | rat | 2.0 | 0.81 | -0.41 | 0.49 | 7025 |
| split2 | rat | 2.0 | 0.41 | -0.96 | -0.64 | 7025 |
| split1 | rat | 6.0 | 0.97 | 0.91 | 0.92 | 7827 |
| split1 | rat | 6.0 | 0.97 | 0.91 | 0.92 | 7827 |
| split1 | rat | 2.0 | 0.96 | 0.90 | 0.90 | 7025 |
| split1 | rat | 2.0 | 0.96 | 0.90 | 0.90 | 7025 |
| split0 | rat | 6.0 | 0.97 | 0.89 | 0.90 | 7827 |
| split0 | rat | 6.0 | 0.96 | 0.90 | 0.81 | 7827 |

**Supplementary Table 5.**
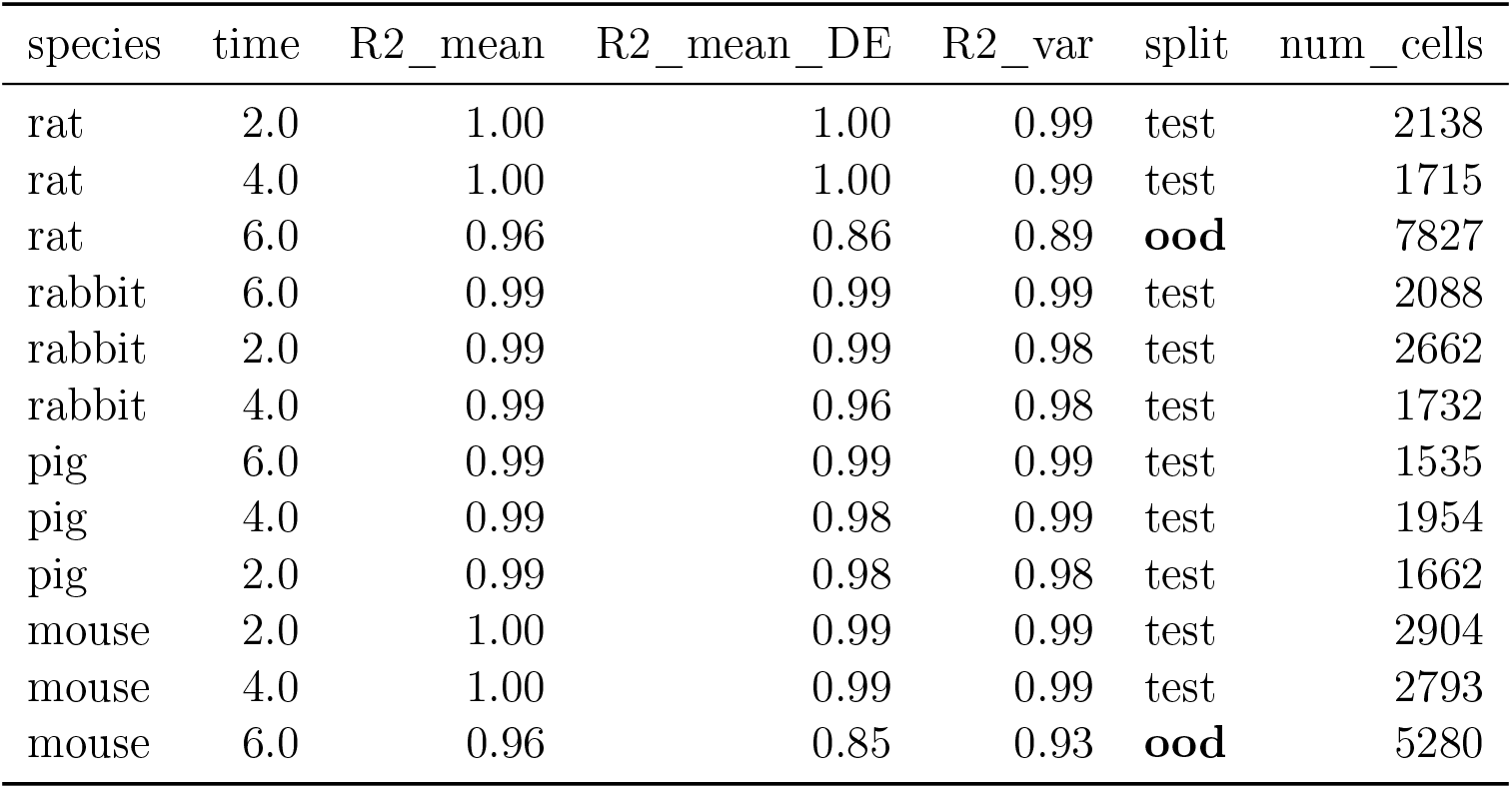
Performance scores for the cross-species dataset.

## Notes

### Summary of Updates

We changed the title of the paper and minor figure improvments.

